# Mechanodetection of neighbor plants elicits adaptive leaf movements through calcium dynamics

**DOI:** 10.1101/2022.01.28.478192

**Authors:** Chrysoula K. Pantazopoulou, Sara Buti, Chi Tam Nguyen, Lisa Oskam, Edward E. Farmer, Kaisa Kajala, Ronald Pierik

## Abstract

Plants detect their neighbors via various cues, including reflected light and touching of leaf tips, which elicit in upward leaf movement (hyponasty). It is currently unknown how touch is sensed and how the signal is transferred from the leaf tip to the petiole base that drives hyponasty. Here, we show that touch-induced hyponasty involves a signal transduction pathway that is distinct from light-mediated hyponasty. We found that mechanostimulation of the leaf tip upon touching causes cytosolic calcium ([Ca^2+^]_cyt_ induction in leaf tip trichomes that spreads towards the petiole. Both perturbation of the calcium response and the absence of trichomes inhibit touch-induced hyponasty. Finally, using plant competition assays, we show that touch-induced hyponasty is adaptive in dense stands of Arabidopsis. We thus establish a novel, adaptive mechanism regulating hyponastic leaf movement in response to mechanostimulation by neighbors in dense vegetation.

## Introduction

Plants growing at high densities compete for resources, including light. Light quality changes are generally known to be received as a neighbor detection cue. The best-established above-ground neighbor detection signal is the reduction of the red (R) to far-red (FR) light ratio (R:FR)^1^ following from waveband-specific absorption (R) and reflection (FR) properties of leaves. Plants respond to reduced R:FR through shade avoidance syndrome (SAS) responses^2,3^, with a central role for PHYTOCHROME INTERACTING FACTORS (PIFs)^4–6^. SAS includes petiole, stem and hypocotyl elongation, apical dominance, early flowering and hyponasty (upward movement of the leaf)^7–9^. Interestingly, different responses are elicited, depending on where low R:FR is perceived: Local FR enrichment of the leaf tip induces differential petiole elongation in the abaxial side of the petiole causing hyponasty, while local FR enrichment of the petiole leads to petiole elongation without hyponasty^10^. Although FR-enrichment by neighbors is a ubiquitously occurring early neighbor detection cue, it is complemented, and sometimes even preceded by, another neighbor detection cue in rosette plant canopies: touching of neighboring leaf tips^11^.

In response to this mechanostimulation of the leaf tip a differential growth response is triggered in the petiole leading to hyponasty, reminiscent of the low R:FR-induced hyponastic response. This hyponastic leaf movement creates a vertical canopy structure that then generates the classic FR light reflection leading to R:FR signaling in plants^11^. The mechanisms involved in detecting and spatially relaying the mechanostimulation from leaf tip to base are currently unknown. Mechanostimulation responses and wounding are primarily regulated by the plant hormone jasmonic acid (JA)^12,13^. Wound responses involve long-distance signaling between the wound and distal tissues that both elicit a rise in JA, and whose JA response patterns are connected via membrane depolarizations processed via Glutamate-receptor like proteins (GLRs)^12,14–17^. Wounding, or mechanostimulation stimulates increases in cytosolic calcium [Ca^2+^]_cyt_. with the GLRs controlling this induction^13,17–22^.

Here, we investigate how leaf tip touching is sensed and signaled over the leaf in order to induced differential growth in the petiole base. We show that the signaling mechanisms of touch-induced hyponasty are fundamentally different from those involved in low R:FR light-mediated hyponasty. In a transcriptome survey we observed strong enrichment of JA- and abscisic-acid (ABA)-associated genes, whereas the canonical auxin pathway was not induced, unlike R:FR-mediated leaf movement. We associated the transcriptome signatures with mechanical stimulation responses, happening specifically in the leaf tip. Using the GFP-based GCaMP3 biosensor, we observed that mechanostimulation of the leaf tip promotes [Ca^2+^]_cyt_ induction and spread towards the petiole in a GLRs-dependent manner. Interestingly, this [Ca^2+^]_cyt_ increase is triggered from the trichomes, the very first tissue to interact between two touching leaves. We show that these are not just the first cells to contact neighbors; they are the sensory structures required to sense and respond to neighbors, since trichome-less mutants lack touch-induced hyponasty.

## Results and discussion

### Distinct signaling pathways regulate touch- and FR-induced hyponasty

In the non-vertically structured canopy of a young Arabidopsis stand, touching of neighboring leaves is the earliest mode of above-ground neighbor detection, eliciting hyponasty^11^. This response entails approximately 20 degrees of upward movement in 24 h exposure against an inert transparent tag mimicking a neighbor leaf (Fig. 1a,b and Supplementary 1a)^11^, and this is further increased after 48 h (Supplementary Fig. 1a). Similar responses are observed in the unrelated species *Nicotiana benthamiana,* (Supplementary Fig. 1b,c), indicating that touch-induced hyponasty is not restricted to *Arabidopsis*. We have recently shown that local reduction of phytochrome activity in the leaf tip through local FR enrichment, on approximately the same position as where leaf-leaf mechanical interactions occur, induces a similar magnitude of hyponasty^10^. This FR-induced hyponasty acts through PHYTOCHROME INTERACTING FACTOR (PIF)4, PIF5 and PIF7 that activate the auxin pathway, at least partially through *YUCCA* (*YUC)8* and *YUC9* gene expression^10^. The resulting elevated auxin is transported and indeed *pin3pin4pin7* triple mutants are not hyponastic in response to FR treatment^10,23^. We therefore, started out by verifying if this pathway is also activated to regulate touch-induced hyponasty. Interestingly, the severe shade avoidance mutants *pif4pif5*, *pif7*, *pin3pin4pin7* and *yuc2yuc5yuc8yuc9*, all showed a wild-type hyponastic response to touch (Fig. 1d-f), whilst being fully irresponsive to FR application (fig. 1g-i). Another auxin biosynthesis mutant *wei8/sav3* and the *yuc8* single mutant also showed a wild type-like touch-induced hyponastic response (Supplementary Fig. 1d,e). Therefore, despite the phenotypic similarity of these two responses, the signaling pathway of touch-induced hyponasty is unique from the core shade avoidance pathway.

**Fig. 1:**
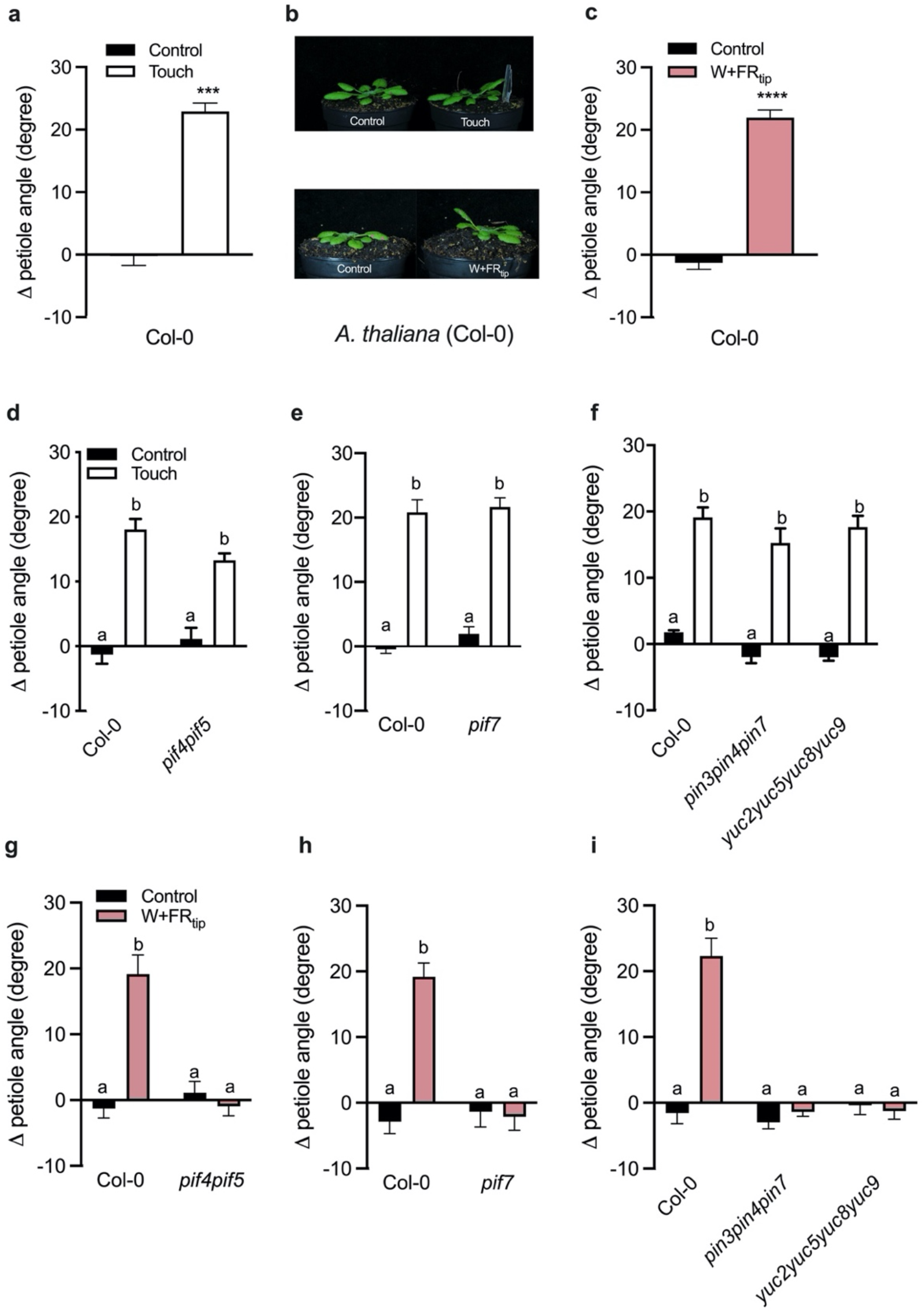
Touch and Local FR -induced hyponasty have a similar phenotypic response but different genetic basis. Differential petiole angle of Col-0 after 24 h (**a**) of touch and (**b**) local-FR treatment. Differential petiole angle of Col-0 compared to (**c**, **d**) *pif4pif5*, (**e, h**) *pif7*, (**f, i**) *pin3pin4pin7* and *yuc2yuc5yuc8yuc9* mutants after 24 h of (**d**-**f**) touch and (**g**-**i**) local FR treatment. The “W+FR_tip_” refers to the local FR treatment of the leaf tip and “Touch” refers to the touch treatment of the leaf tip with an inert transparent tag. Data represent mean ± SE (n= 8-10). Different letters indicate significant difference (two-way ANOVA with Tukey’s post hoc test; P<0.05).

### Transcriptome analysis reveals specific regulation in petiole versus leaf tip

To unveil the mechanisms of touch-induced hyponasty in depth, we analyzed transcriptome data (using Affymetrix *Arabidopsis* Gene 1.1 ST arrays), comparing the site of perception (leaf tip) and the site of action (petiole base) (Fig. 2a) under control and touch conditions. We found similar numbers of up- and downregulated genes between the two tissues (Fig. 2b), but interestingly there was nearly no overlap in differentially expressed genes (DEGs) (Fig. 2c), indicating that different parts of the leaf have distinct transcriptional responses to touch. We then compared the touch-induced hyponasty transcriptome data with the previously published local FR-induced hyponasty transcriptome data that were collected from the same leaf tissues and under identical conditions in the same run of experiments^10^. Touch induced only a minor number of DEGs (less than 100) compared to the low R:FR treatment (over 700 DEGs) in the leaf tip, and also in the petiole base the number of DEGs in low R:FR treatment is almost 5 times higher than in the touch treatment (Supplementary Fig. 2a). Amongst the leaf tip DEGs only 35 genes overlapped between touch and low R:FR treatments. Although this is approximately half of the touch-induced genes, the genes themselves (excel file 1; are not typically associated with the canonical shade avoidance machinery, but rather with abscisic acid (ABA) response, jasmonic acid (JA) metabolism and cell wall homeostasis (Fig. 2d, Supplementary Fig. 2d). Gene ontology (GO) enrichment analysis on the full touch-induced DEG profiles indeed revealed a high representation of JA and ABA-related processes in the leaf tip and petiole base respectively (Fig. 2d). Hence, we tested the role of JA on touch-induced leaf hyponasty. Application of high exogenous MeJA (methyljasmonate) to the leaf blade, inducer of JA-signaling pathway, did not inhibit touch-induced hyponasty (Supplementary Fig. 3a). Also, the touch response was not different from the wild type in the JA biosynthesis mutants *lox2*, *loxQ* (*lox2lox3lox4lox6*) and *aos* and in the JA receptor mutant *coi1-34* (Supplementary Fig. 3 b,c). MYC transcription factors act downstream of JA activation and although the *myc* single mutants were similar to the wild type, the *myc2myc3myc4* triple mutant had a mildly reduced response (Supplementary Fig. 3d,e). Since the transcriptome analysis also revealed ABA signatures in the petiole base, we first applied ABA to the petiole, which reduced hyponasty only at the highest concentration (Supplementary Fig. 3f), probably due to an overall inhibition of petiole growth (Supplementary Fig. 3g). Mutants for ABA biosynthesis (*aba2-1*, *aba3-1*), ABA perception (*pyr1pyl1pyl2pyl4*, referred to as *abaQ*) and ABA signaling (*areb1areb2abf3abf1,* referred to as *arebQ*) all responded similar to Col-0 wild type (Supplementary Fig. 3h-j). Summarizing, ABA and MeJA application can limit petiole growth and thereby hyponasty, but a wide variety of mutants for the JA and ABA pathways suggest no major role for these two hormones in touch-induced hyponasty.

**Fig. 2:**
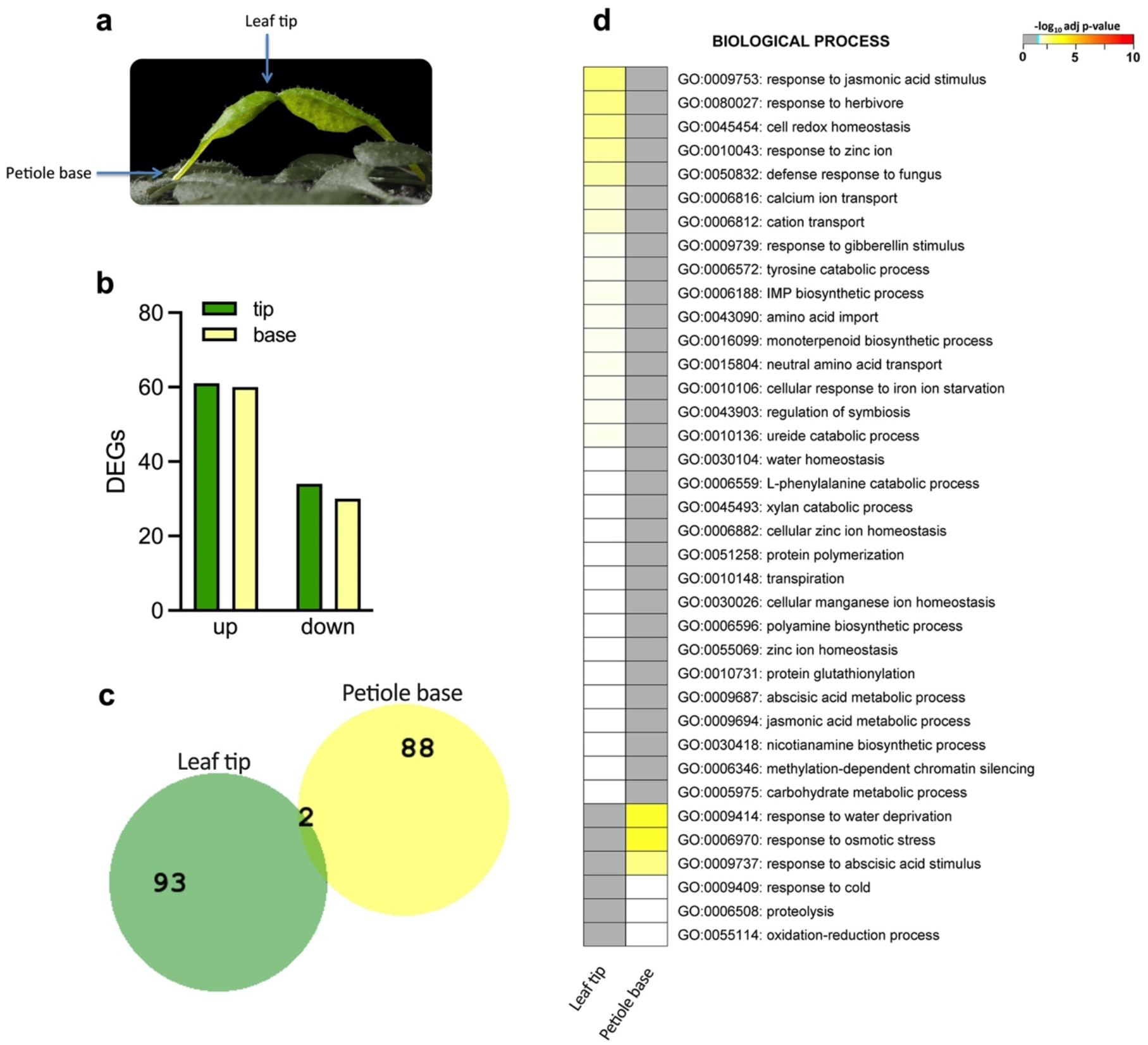
Comparative analysis of touch-induced hyponasty in the leaf tip and the petiole base. (**a**) Leaf tip and petiole base tissues were harvested, in the experiment touch was induced by an inert transparent tag. (**b**) Number of differentially expressed genes (DEGs) in the lamina tip and the petiole base. (**c**) Comparison of DEGs in leaf-tip (“Leaf tip”) and petiole base (“Petiole base”) in response to touch (adj. P-value < 0.05). (**d**) GO enrichment analysis of the DEGs in the leaf tip (“Tip”) and the petiole base (“Base”).

### Touch-induced hyponasty is mediated via changes in cytosolic [Ca^2+^]

Mechanostimulation responses are highly associated with cytosolic calcium dynamics ([Ca^2+^]_cyt_) following mechanical perturbation in many species, including *Arabidopsis* ^17,21,24–26^. Specifically, mechanical perturbation has been found to increase intracellular calcium by triggering temporary changes in the cytosolic calcium concentration^18,22,27,28^. Indeed, our transcriptome data showed GO enrichment of calcium ion transport in the leaf tip upon touch (Fig. 2d). To verify if [Ca^2+^]_cyt_ is affected during touch-induced hyponasty, we used the GFP fluorescence-based cytosolic calcium biosensor *UBQ10pro::GCaMP3* that allows detection of [Ca^2+^]_cyt_ fluxes in the leaf^17,21,25^. Upon gently touching the 5^th^ youngest leaf of 4-week old *Arabidopsis* we recorded the GCaMP3 fluorescence dynamics. We measured GCaMP3 fluorescence with a microscope positioned above the leaf, in the leaf tip (position 1), in two subsequent positions of the primary vein of the leaf (positions 2 and 3), the leaf blade-petiole junction (position 4) and the middle and base of the petiole (positions 5 and 6 respectively) (Fig. 3a,b). We observed that GCaMP3 fluorescence started to increase in 4 min (250 sec) upon gentle touch (Video 1) in all the positions except position 6 (Fig. 3c; table 1). This is substantially slower than what has been reported in wound or other touch responses, where changes were recorded as early as 30 seconds after stimulation^17,21,26,29^. Clearly, the mechanical force of stimulation in the experiments here is much lower than during severe wounding or even the rapid leaf closure of carnivorous Venus flytrap upon capture of the prey^30,31^. Possibly the degree or force of the mechanical stress could affect the rapidity of induction of [Ca^2+^]_cyt_ flux. Touching of the leaf tip (position 1) led to a peak GCaMP3 signal at 13 min (780 sec) upon touch in the leaf tip itself (position 1), followed by a gradual decrease of the fluorescence signal. The calcium wave progressed from the leaf tip through the leaf and into the petiole, but fluorescence intensity decreased with distance from the site of mechanostimulation (Fig. 3c; table 1). The GCaMP3 fluorescence in the petiole (positions 5 and 6) was rather weak and this may indicate only very weak [Ca^2+^]_cyt_ signal reaching the base, or a preferential localization to the abaxial side that cannot be imaged in our microscope configuration. To verify that the observed [Ca^2+^]_cyt_ dynamics were required for touch-induced hyponasty, we used the Ca^2+^ channel inhibitor LaCl_3_ on the touched leaf and found that touch-induced hyponasty was strongly reduced (Supplementary Fig. 4a). Although LaCl_3_ treatment could give pleiotropic effects, we observed that LaCl_3_ did not affect the hyponastic response to local FR treatment at all, suggesting the effect of the inhibitor to be specific to touch-induced hyponasty (Supplementary Fig. 4b).

**Fig. 3:**
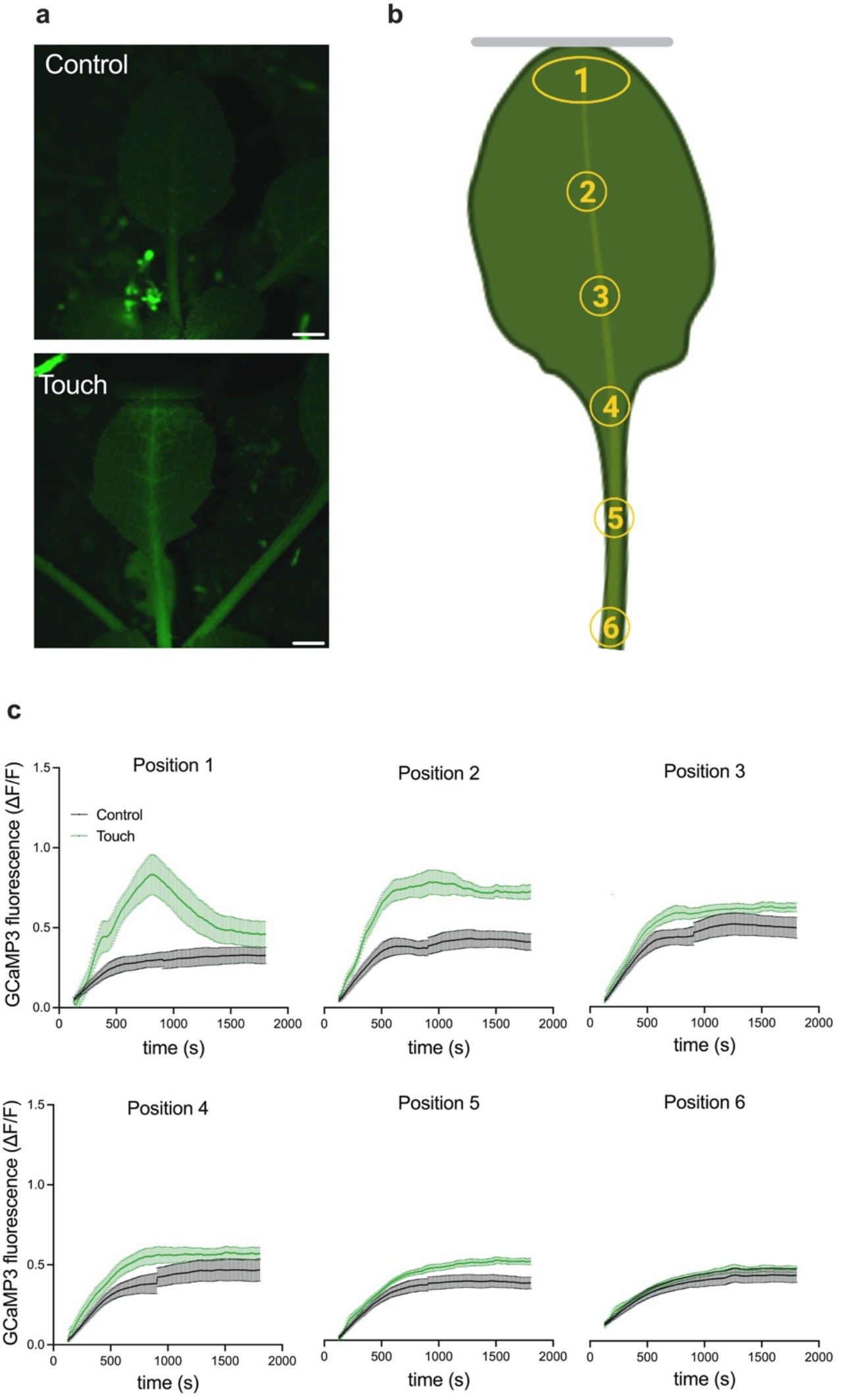
Touch-induced hyponasty causes cytosolic Ca^2+^ increase in the touched leaf. (**a**) Fluorescence induction in the leaf lamina after 20 min of untouched (Control) or touch treatment (Touch) using the fluorescent cytosolic calcium biosensor *UBQ10pro::GCaMP3*. (**b**) Six different positions were used to measure the GCaMP3 fluorescence in leaf upon control or touch treatment. Grey line represents the transparent tag. (**c**) Time course of GCaMP3 fluorescence intensity in leaf tip (position 1), primary vein of the lamina (position 2 and 3), lamina-petiole junction (position 4), in the middle of the adaxial site of the petiole (position 5) and in the adaxial site of the petiole base (position 6) upon control (black line) and touch (green line) treatment. Touch was induced by a transparent tag. Data represent mean ± SE; n = 8-10. Scale bar correspond to 1 mm. Panel **b**was created with Biorender.

Clade 3 Glutamate receptor-like proteins (GLRs) have been associated with propagation of long-distance electrical signals via [Ca^2+^]_cyt_ after mechanical stress^12,17,21,22^. Nguyen et al. (2018)^17^ showed that GLR3.1, GLR3.3, GLR3.6 are localized in the vasculature and a calcium signal is severely attenuated in the *glr3.1glr3.3* and *glr3.3glr3.6* double mutants. Since the calcium waves observed here (Fig. 3c) start from the leaf tip and move to the petiole along the primary vein, we investigated the involvement of these GLRs in touch-induced hyponasty. Indeed, the *glr3.3glr3.6* double mutant showed a slight reduction to touch-induced hyponasty (Fig. 4a). Given the rather weak effect, we also created a triple knockout mutant *glr3.1glr3.glr3.6* and found that its touch-induced hyponasty was strongly attenuated (Fig. 4b,c), indicating that clade 3 GLRs may redundantly regulate touch-induced hyponasty. This triple mutant has wild-type-like hyponasty in response to local FR treatment (Supplementary Fig. 4c), reinforcing the specificity of the [Ca^2+^]_cyt_-associated regulatory route for mechanostimulation.

**Fig. 4:**
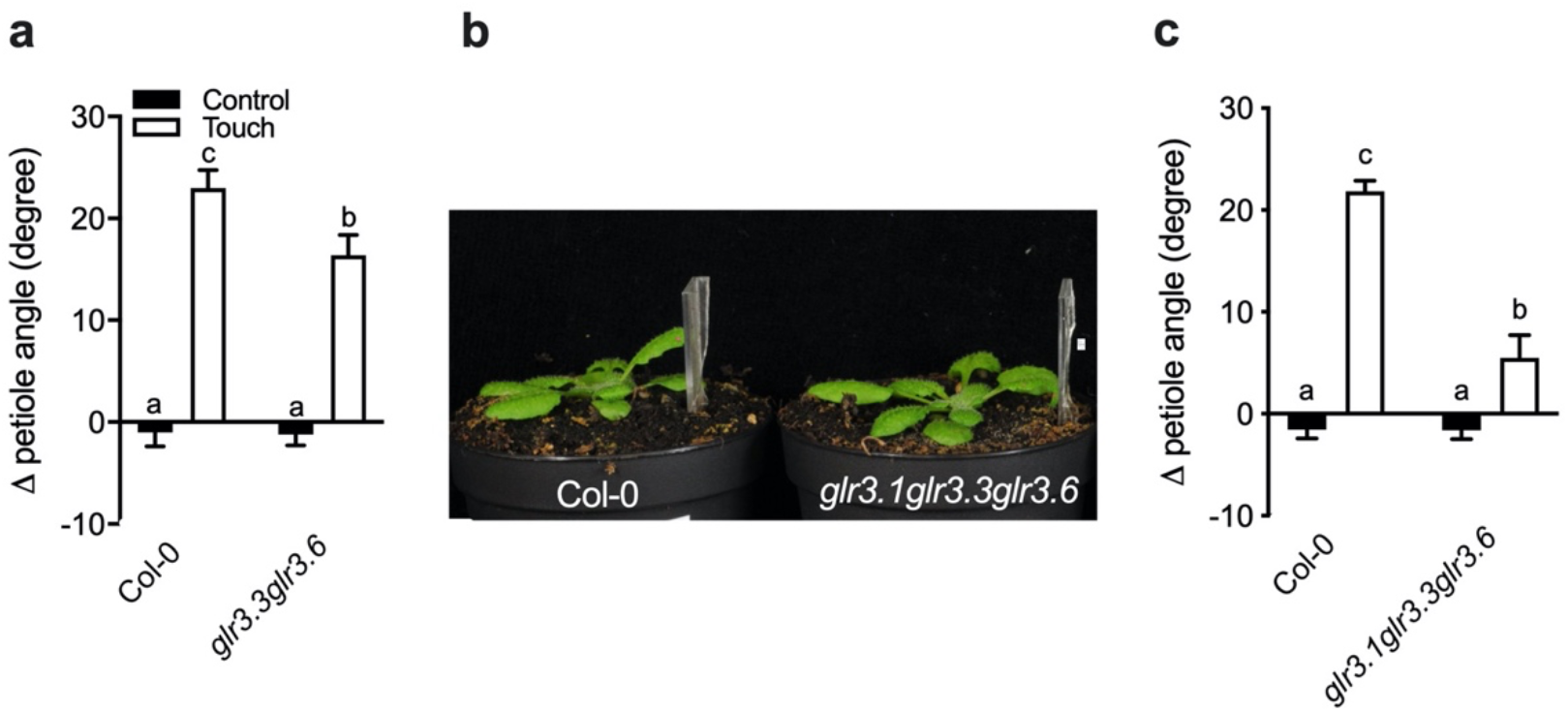
GLRs are required for the full touch-induced hyponasty response. (**a**) Differential petiole angle of Col-0 compared to *glr3.3glr3.6* double mutant after 24 h of touch treatment. (**b, c**) The touch response of the *glr3.2glr3.3glr3.6* triple mutant is significantly reduced compared to Col-0 after 24 h of touch treatment. Touch was induced by a transparent tag. Data represent mean ± SE; n = 10-16. Different letters indicate significant differences (two-way ANOVA with Tukey’s post hoc test; P < 0.05).

### Trichomes are required to sense and respond to touch

Trichomes on the leaf edge are the first cells to physically interact with neighboring leaves when they grow towards each other in dense stands. Previous work established that trichomes can trigger calcium oscillations in the trichome stalk and the trichome base cells surrounding the trichomes upon mechanical perturbation^32^. We monitored GCaMP3 fluorescence at three trichome positions upon gentle trichome touching: the trichome stalk (position 1, Fig. 5a), the trichome base cells (position 2) and the epidermal cells around the trichome (position 3) (Supplementary Fig. 5a-c). We also monitored how far the calcium can spread upon trichome-touch by recording an area adjacent to the neighboring trichome (position 4, Supplementary Fig. 5a,d). A strong induction of the GCaMP3 fluorescence was detected in the trichome stalk (position 1) 30 sec after very gently touching the trichome with a toothpick (Video 2;), which rapidly spread to the base cells (position 2) and the epidermal cells around the trichome (position 3) (Fig. 4a,b, Supplementary Fig 5b,c). Interestingly, GCaMP3 fluorescence was detected near the neighboring trichome (position 4) suggesting that the [Ca^2+^]_cyt_ induction was strong enough to extend laterally to the untouched trichome area (Supplementary Fig. 5d). Our findings indicate that that gentle touching of just the trichomes can lead to calcium induction and spreading across the leaf blade, consistent with our observations that leaf tip touch can elicit a [Ca^2+^]_cyt_ wave along the leaf blade towards the petiole. Since trichomes are the first cells to touch neighboring leaves, and they can generate [Ca^2+^]_cyt_ changes that progress through the leaf, we hypothesized that trichomes could be the mechanosensitive sensors of neighbor plants. To verify the role of trichomes in touch-induced hyponasty, we compared the Col-0 accession with a glabrous (non-trichome-forming) *Arabidopsis* accession, known as 9354 (N28001)^33^. Accession 9354 indeed had a strongly reduced touch-induced hyponasty, as compared to Col-0 (Fig. 5c,d). Similar findings were also obtained using other glabrous *Arabidopsis* accessions (Fran-3, Wil-2, Br-0) upon touch treatment (Supplementary Fig. 5e). The *TTG1*^34^ and *GL1*^35^ genes are positive regulators of *Arabidopsis* trichomes, and their respective mutants, *ttg1* and *gl1*, do not form trichomes. These two mutants displayed severely reduced touch-induced hyponasty as well (Fig. 5e). At the same time, all the glabrous genotypes had wild-type-like local FR-induced hyponasty (Supplementary Fig. 5f,g), indicating that trichomes are specifically involved in the touch-induced hyponasty.

**Fig. 5:**
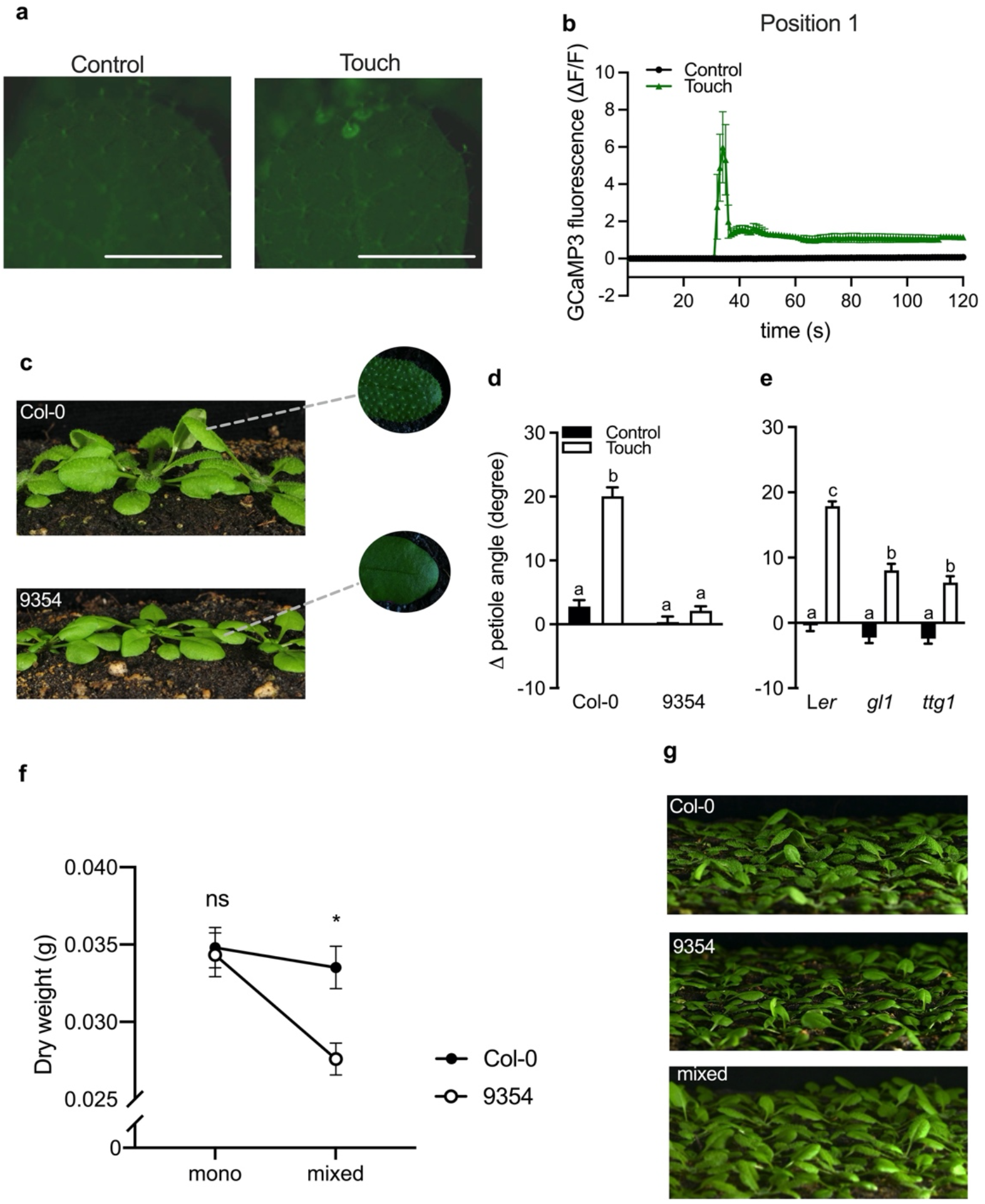
Trichomes are required for the touch-induced hyponasty. (**a**) GCaMP3 fluorescence in trichomes after seconds of untouched (Control) or touch treatment (Touch), using the fluorescent cytosolic calcium biosensor *UBQ10pro::GCaMP3*. (**b**) Time course of GCaMP3 fluorescence in trichome stalk of Col-0 upon control or touch treatment. (**c**) Hyponastic responses are seen following leaf touching of densely grown Col-0 (with trichomes) but not in 9354 (without trichomes) plants. (**d**, **e**) Differential petiole angle of several genotypes that lack trichomes compared to their wild-type after 24 h of touch treatment: (**d**) Col-0 and 9354 and (**e**) L*er, ttg1* and *gl1*. Touch was induced by a transparent tag. (**f**) Dry weight of Col-0 and 9354 growing in monoculture canopies and in a mixed canopy (Col-0 together with 9354). (**g**) Pictures illustrating (from the top to the bottom) the monoculture canopy of Col-0 and 9354 and the mixed canopy of both genotypes together. Data represent mean ± SE; n = 8-14. Different letters or asterisks indicate significant differences (two-way ANOVA with Tukey’s post hoc test or t-test; P < 0.05). Scale bar correspond to 1 mm.

After establishing the trichome - [Ca^2+^]_cyt_ mechanism regulating touch-induced leaf movement, an important question remained: Is the touch-induced hyponasty of quantitative importance to plant performance in dense stands? To answer this question we grew single plants, as well as monoculture stands and 1:1 mixed stands of the accessions Col-0 and 9354. We kept their root systems separated to prevent belowground competition, and measured shoot dry weight as a proxy of plant performance (Fig. 5f,g). Importantly, Col-0 and 9354 showed identical growth when grown individually or when grown in crowded monocultures of each of the genotypes, but in the mixture where they interacted above-ground, Col-0 outcompeted 9354 (Fig. 5f,g). We, therefore, propose that the trichome-based detection of neighboring leaves and the corresponding hyponasty, promote plant performance in competition for light. Suboptimal ability to do so, as in the 9354 accession, leads to reduced competitive performance. Future studies may investigate the targets of trichome-derived [Ca^2+^]_cyt_ for differential growth between the abaxial and adaxial side of the petiole base, presumable involving regulators other than those involved in R:FR responses. Since hyponasty enhances the vertical element of a rosette canopy structure that then stimulates FR reflection to neighbors^10^, the novel signaling route through trichomes and [Ca^2+^]_cyt_ dynamics precedes the better-known photoreceptor-driven pathways that are activated once a notable vertical canopy structure has been created.

## Methods

### Plant materials, growth and measurements

Genotypes used in this study that are in the Col-0 background are: *pif4-101 pif5-1*^36^, *pif7-1*^37^, *pif4-101pif5-1pif7-1*^4^, *abaQ, aba2-1, aba3-1*^38^, *arebQ*^39^, *myc2*^40^, *myc3, myc4, myc2myc3myc4* ^41^, *glr3.3aglr3.6a* ^12^, *glr3.1aglr3.3aglr3.6a* were generated by crossing *glr3.3aglr3.6a* and *glr3.1a16, aos*^42^, *coi1-34*^43^, *lox2* ^15^, *loxQ*^44^, *wei8*^45^, *pin3-3pin4pin7* ^46^, *yuc2yuc5yuc8yuc9*^47^, *yuc8*^48^. Genotypes used in this study in the L*er* background are: *gl1* and *ttg1*^49^. We also used *Arabidopsis* accessions without trichomes 9354 (N28001), Fran-3 (N75673), Wil-2 (N1596), Br-0 (N22628) and *N. benthamiana*. Seeds were sown on Primasta^®^ soil and stratified for 3 days (dark, 4°C), before transferring to short day (9 h light / 15 h dark) growth rooms (130-135 μmol m^−2^ s^−1^ PAR, R:FR 2.3, 20°C, 70% RH). After 11 d, seedlings were transplanted to 70 ml pots for all experiments, except for the competition assays. For touch experiments an inert, transparent tag (polycyclical olefin) was placed in the soil, next to the fifth youngest leaf of the 28-days old plant to mimic leaf-leaf touching. Petiole angles to the horizontal of the fifth-youngest leaf that just touched the transparent tag were determined with image-J software (https://imagej.nih.gov/ij/download.html) from pictures taken just before (t = 0 h) and after treatment (t = 24 h). All experiments started at 10:00 in the morning (ZT = 2 h). For competition assays 11 days old seedlings were transplanted to tree trays with individual pots (to avoid any root competition) to create dense stands of 7 x 7 plants (pots = 19 ml and distance between the plants 2.2 cm). Col-0 and 9354 plants were transplanted to monoculture (Col-0 or 9354) and mixed (Col-0 and 9354 in a 1:1 checkerboard grid^50^) canopies plots. Plants in the outer rows of the canopy served to minimize edge effects and only plants in the middle of the canopy were harvested when the canopy plots were 35 days old. Shoots were dried in an oven at 70°C for three days and shoot dry weight of each individual plants was recorded.

### Far-red (FR) light treatments

Supplemental FR light treatments were performed using a localized FR beam focused on the leaf tip (see Pantazopoulou et al., 2017^10^ for details) and abbreviated as W+FR_tip_. All growth conditions were standard as mentioned above, except that locally on the leaf tip R:FR dropped from 2.3 to 0.05. The R:FR treatment had no effect on the photosynthetically active radiation.

### Pharmacological experiments

Plants were treated with different concentrations of the hormones MeJA (Van Meeuwen Chemicals BV, NL), or ABA (Sigma-Aldrich, USA). MeJA was given to the leaf blade, whereas ABA was applied to the petiole. All solutions, including the mock treatments contained 0.1% DMSO and 0.1 % Tween. The solutions were freshly made and they were applied right before and 5 h after the touch treatment. LaCl_3_ (Sigma-Aldrich, USA) treatment was done (2mM of LaCl_3_ application to whole leaf) 24h before the touch or W+FR_tip_ treatment. The solution was freshly made with water and 0.1 % Tween.

### Transcriptome data analysis

The transcriptome data on touch treatment were collected in a larger experiment that also included local FR treatments, and we published the FR transcriptome data in Pantazopoulou et al., 2017^10^. The harvesting, extraction, and processing as well statistical analyses are essentially as previously published^10^. In brief, the leaf tip and petiole base from wild-type plants (Col-0) were harvested after 5 h of touch treatment. 15 petiole bases and 15 leaf tips were pooled for each sample for RNA extraction [three biological replicates (independent experiments) per tissue per treatment, collected from three independent experiments]. Affymetrix 1.1 ST *Arabidopsis* arrays were used to hybridize the samples via a commercial provider (Aros, Aarhus, Denmark). The raw data were normalized for signal intensity to remove background noise. The quality check of the data was performed using Bioconductor (packages “oligo” and “pd.aragene.1.1.st”) in R software. Differential expression analysis was carried out using the Bioconductor ‘’Limma’’ package in R software. Genes with adjusted p-value < 0,05 were considered as differentially expressed. Gene ontology (GO) analysis was done with GeneCodis (http://genecodis.cnb.csic.es). Clustering was based on the positive and negative logFC for each set. The data are available in the National Center for Biotechnology Gene Expression Omnibus database (https://www.ncbi.nlm.nih.gov/geo/query/acc.cgi; accession no. GSE98643).

### GCaMP3 fluorescence visualization and quantification

The GCaMP3 fluorescence is quantified via the ΔF/F ratio. ΔF/F = (F-F_0_)/F_0_, where F is the GCaMP3 fluorescence of a given time point during touch while F_0_ is the averaged based line in the ROIs in the first 2 min before touch treatment. The GCaMP3 fluorescence calculation for the leaf was performed in each selected leaf position for every ten seconds, while in the trichomes for every second. Video recordings were made with a 1.5x objective on an SMZ18 stereomicroscope (Nikon Instruments Europe BV, Amsterdam, Netherlands) equipped with an ORCA-Flash4.0 (C11440) camera (Hamamatsu, Solothurn, Switzerland) and eGFP emission/excitation filter set (AHF Analysentechnik AG, Tübingen, Germany). Light was supplied to the stereomicroscope using fibre optics. Video with a resolution of 512 x 512 pixels were acquired using NIS-Elements software (Nikon) with 1 frame s-1 frequency. Recordings were carried out in the dark at 22°C.

### Statistical analysis

Data were analyzed with one or two-way ANOVA followed by Tukey’s HSD test using GraphPad and Rstudio.

## Supporting information

excel file 1_DEGs

Video 1 -leaf tip

Video 2 -trichome

## ACKNOWLEDGEMENTS

We thank the Plant Ecophysiology group (UU) for help with tissue harvests for the transcriptome experiments, Yorrit van de Kaa for preparing the plant material for the experiments, Debatosh Das and Ronnie de Jong for help with bioinformatics analysis. This work was funded by Netherlands Organization for Scientific Research: Vidi Grant 86512.003 (R.P. and C.K.P.) and ALW open grant ALWOP.509 (C.K.P. and K.K.).

## AUTHOR CONTRIBUTIONS

Chrysoula K. Pantazopoulou and Ronald Pierik designed research, with additional input from Edward E. Farmer; Chrysoula K. Pantazopoulou, Sara Buti Chitam Nguyen and Lisa Oskam performed research; Chrysoula K. Pantazopoulou, Sara Buti, Chitam Nguyen and Kaisa Kajala analysed data; Chrysoula K. Pantazopoulou wrote the manuscript draft; Edward E. Farmer, Kaisa Kajala and Ronald Pierik revised the manuscript.

## Notes

### Competing Interest Statement

The authors have declared no competing interest.

