## Supplemental figures and table for "Mechanodetection of neighbor plants elicits adaptive leaf movements through calcium dynamics"

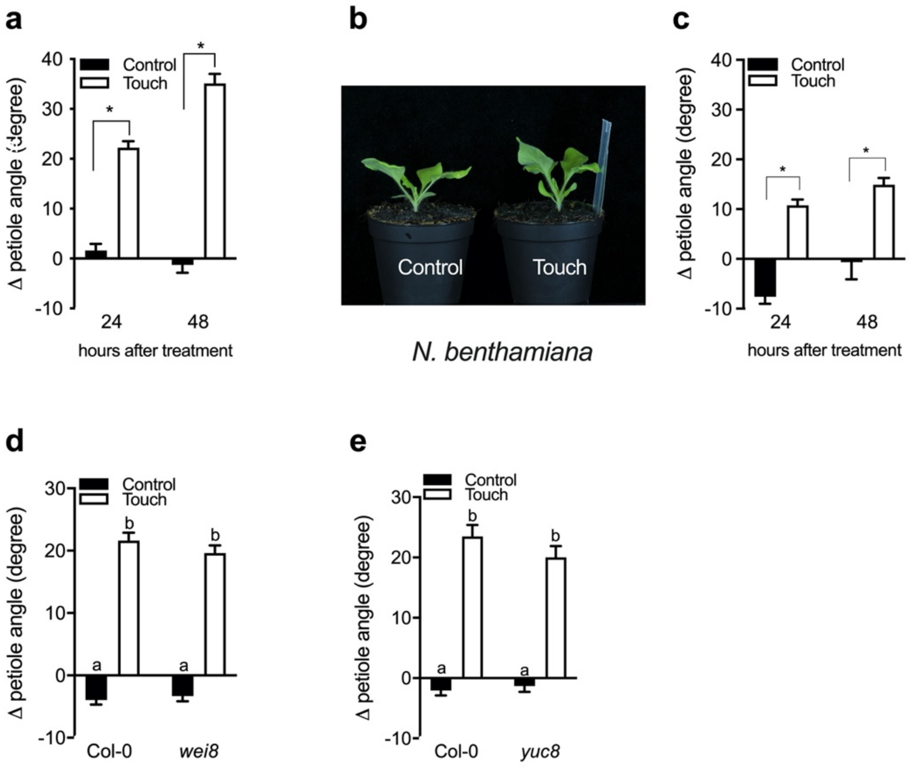


**Supplementary Figure 1**: Touch-induced hyponasty in *A. thaliana* and *N. benthamiana*, as well as in *A. thaliana* auxin biosynthesis mutants. The differential petiole angle of touch-induced hyponasty in (**a**) *A. thaliana* Col-0 and (**c**) *N. benthamiana* after 24 h and 48 h. Representative photo of (**b**) *N. benthamiana* leaves touching transparent tags after 24 h. The differential petiole angle of touch-induced hyponasty in Col-0 compared to (**d**) *wei8* and (**e**) *yuc8* mutants 24h after touch treatment. Touch was induced by an inert transparent tag. Data represent mean ± SE; n = 8-10. Statistically significant differences are indicated with asterisk, paired Student's t test ($P < 0.05$).

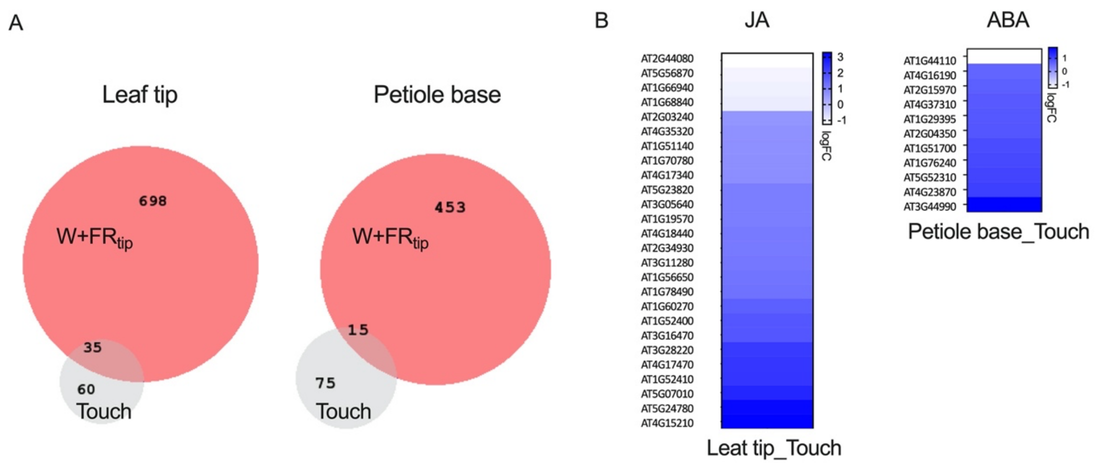


**Supplementary Figure 2**: Different molecular pathways regulate the touch- and FR-induced hyponasty. (**a**) Venn diagrams illustrate the differentially expressed genes (DEGs) common to the touch and local-FR treatment in the leaf tip (the first venn diagram) and petiole base (the second Venn diagram). (**b**) Heatmap representation of the change in expression level of DEGs that are associated with JA and ABA in the leaf tip (“Leaf tip_Touch”) and petiole base (“Petiole base_Touch”) respectively in response to touch.

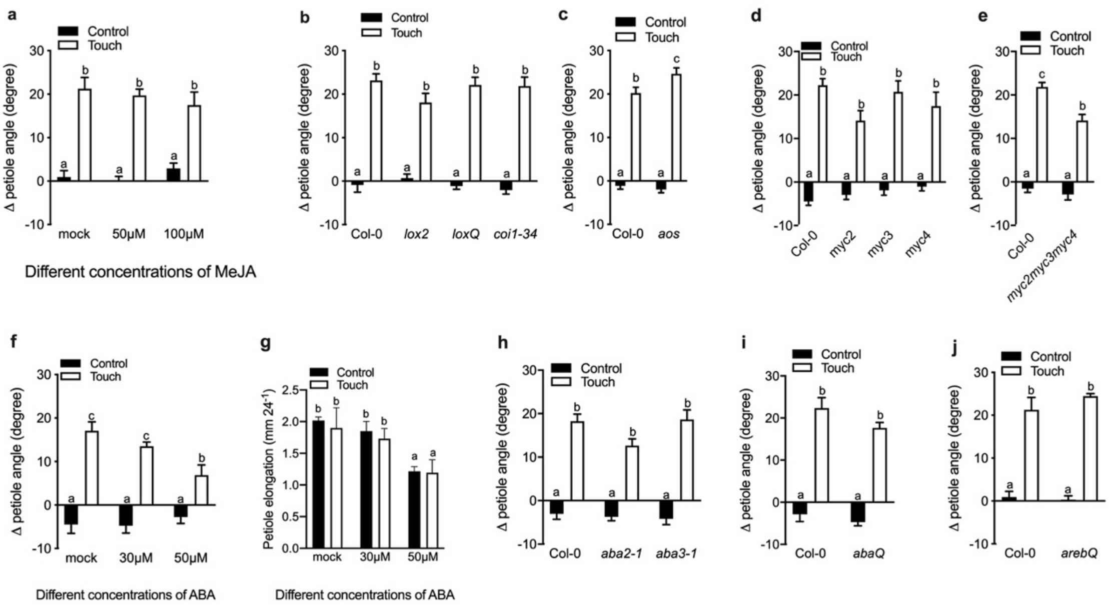


**Supplementary Figure 3**: Touch-induced hyponasty is not specifically regulated by ABA or JA. Differential petiole angle after exogenous application of different (**a**) MeJA (50 µM, 100 µM and 200 µM) concentrations after 24 h of touch treatment. Differential petiole angle of Col-0 compared to (**b**) *lox2*, *loxQ*, *Coi-34*, (**c**) *aos* (**d**) *myc2*, *myc3*, *myc4*, and (**e**) *myc2myc3myc4* mutants after 24 h of touch treatment. (**f**) Differential petiole angle and (**g**) petiole elongation after exogenous application of different ABA concentrations (30µM and 50µM) after 24h of touch treatment. Differential petiole angle of Col-0 compared to (**h**) *aba2-1* and *aba3-1*, (**I**) *abaQ* and (**j**) *arebQ* mutants after 24 h of touch treatment. Touch was induced by a transparent tag. Data represent mean ± SE; n = 6 -16. Different letters indicate significant differences (two-way ANOVA with Tukey's post hoc test; $P < 0.05$).

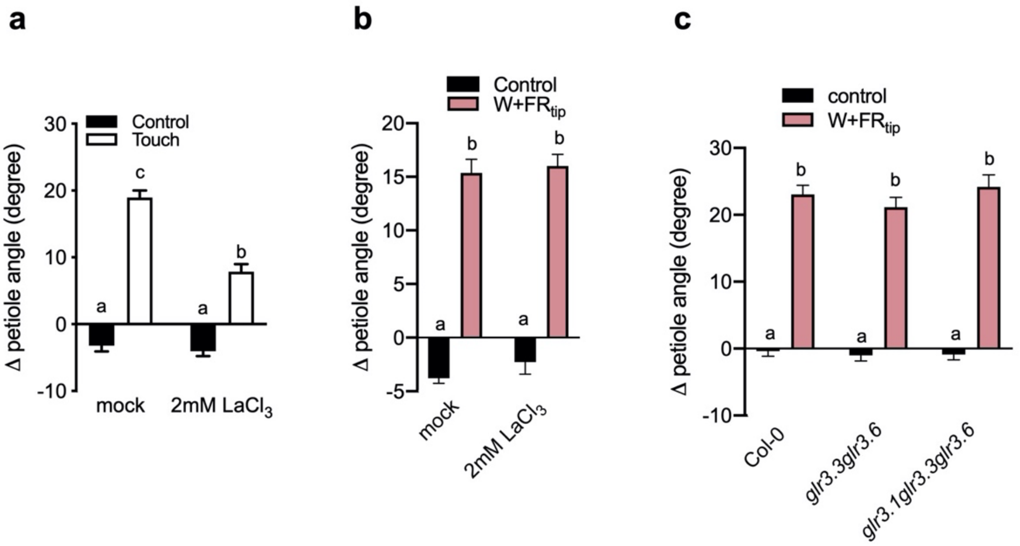


**Supplementary Figure 4**: The *glr* mutants and $LaCl_3$-treated plants have normal hyponasty response to local FR treatment. Differential petiole angle of Col-0 after 24h of (**a**) touch (“Touch”) or (**b**) local FR (“W+$FR_{tip}$”) treatment. (**c**) Differential petiole angle of Col-0 compared to *glr3.3aglr3.6a* and *glr3.1glr3.3glr3.6,* after 24 h of local FR treatment (W+$FR_{tip}$). Touch was induced by a transparent tag. Data represent mean ± SE; n = 7-14. Different letters indicate significant differences (two-way ANOVA with Tukey’s post hoc test; $P < 0.05$).

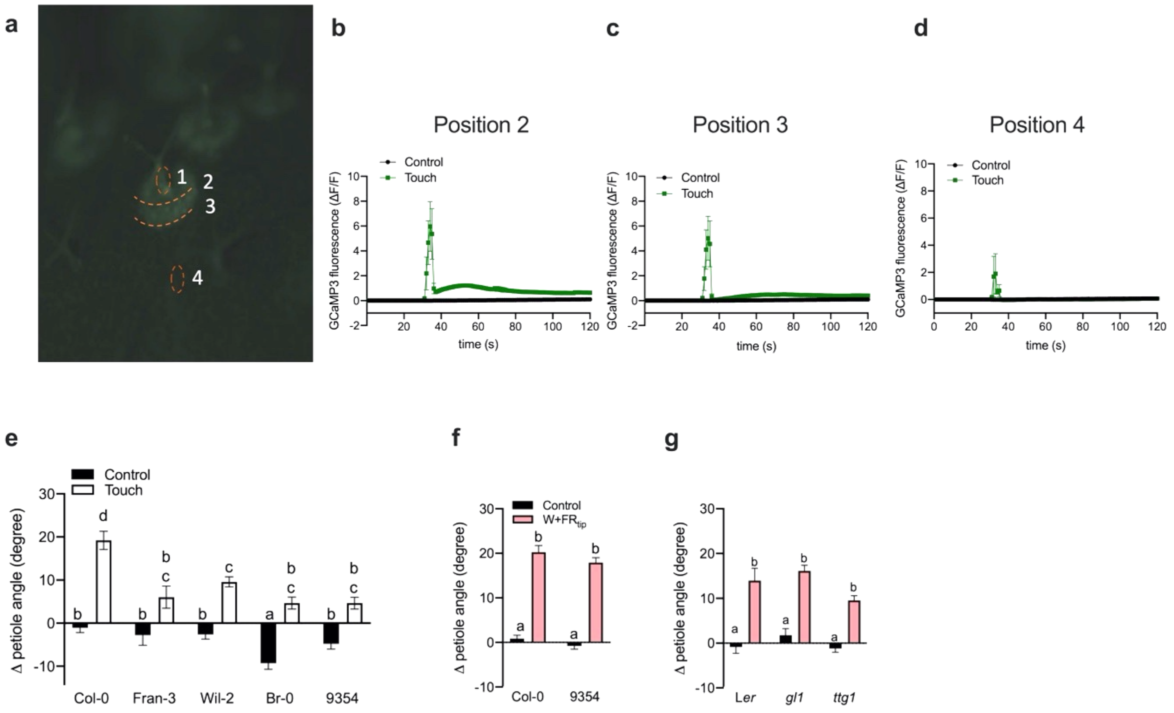


**Supplementary Figure 5**: Trichomes play a key role in the touch-induced hyponasty but not in the local FR-induced hyponasty. (**a**) Four different positions were used to measure the GCaMP3 fluorescence within the trichome upon control or touch treatment, using the fluorescent cytosolic calcium biosensor *UBQ10p::GCaMP3*. (**b**-**d**) Time course of GCaMP3 fluorescence in skirt cells (position 2), surrounding trichome cells (position 3) and close to a neighbor trichome (position 4) upon control (black line) and touch (green line) treatment. For position 1, see Fig. 5b. Touch was induced by a transparent tag. (**e**)Differential petiole angle of Arabidopsis accessions without trichomes (Fran-3, Wil-2, Br-0, 9354) compared to an accession with trichomes (Col-0) after 24 h of touch treatment. (**f**, **g**) Differential petiole angle of (**f**) 9354 and (**g**) *gl1* and *ttg1* compared to their wild types, after 24 h of local FR treatment ($W+FR_{tip}$). Data represent mean ± SE; n = 7-12. Different letters indicate significant differences (two-way ANOVA with Tukey's post hoc test; $P < 0.05$).

**Table 1**: Statistical analysis of Fig. 3**c**. $P < 0.05$ indicate statistically significant differences (Repeated measures ANOVA), while $P > 0.05$ indicate no statistically significant differences.

| time | 0 - 500 sec | | | 500 - 1000 sec | | |
|---|---|---|---|---|---|---|
| | treatment | time | Treatment:Time | treatment | time | Treatment:Time |
| Position 1 | P=0.115 | p<0.0001 | p<0.0001 | p<0.01 | p<0.0001 | p=0.0017 |
| position 2 | p<0.01 | p<0.0001 | p<0.0001 | p=0.0005 | p<0.0001 | p=0.0064 |
| position 3 | p=0.2153 | p<0.0001 | p<0.0002 | p=0.068 | p<0.0001 | p=0.1629 |
| position 4 | p=0.1952 | p<0.0001 | p<0.01 | p=0.0693 | p<0.0001 | p<0.0001 |
| position 5 | p=0.3767 | p<0.0001 | p<0.01 | p=0.0413 | p<0.0001 | p<0.0001 |
| position 6 | P=0.9171 | p<0.0001 | P=0.1435 | P=0.0504 | p<0.0001 | p<0.0001 |

**Video 1**: Leaf tip touch increaes $[Ca^{2+}]_{cyt}$ as reported with the cytosolic calcium biosensor *UBQ10p::GCaMP3*. The video is 85 x real time, time stamp indicates minutes and seconds of real time. Touch was induced by a transparent tag.

**Video 2**: Touching trichomes at the leaf tip stimulates $[Ca^{2+}]_{cyt}$ as reported with the cytosolic calcium biosensor *UBQ10p::GCaMP3*. The video is 6 x real time, time stamp indicates minutes and seconds of real time . Touch was induced by a toothpick.
